# Privacy-preserving impact evaluation using Difference-in-Differences

**DOI:** 10.1101/2023.12.05.570107

**Authors:** Manuel Huth, Carolina Alvarez Garavito, Lea Seep, Laia Cirera, Francisco Saúte, Elisa Sicuri, Jan Hasenauer

## Abstract

Difference-in-Differences (DID) is a widely used tool for causal impact evaluation but is constrained by data privacy regulations when applied to sensitive personal information, such as individual-level performance records or healthcare data, that must not be shared with data analysts. Obtaining consent can reduce sample sizes or exclude treated/untreated groups, diminishing statistical power or making estimation impossible. Federated Learning, which shares aggregated statistics to ensure privacy, can address these concerns, but advanced federated DID software packages remain scarce. We derived and developed a federated version of the Callaway and Sant’Anna DID, implemented within the DataSHIELD platform. Our package adheres to DataSHIELD’s security measures and adds extra protections, enhancing data privacy and confidentiality. It reproduces point estimates, asymptotic standard errors, and bootstrapped standard errors equivalent to the non-federated implementation. We demonstrate this functionality on simulated data and real-world data from a malaria intervention in Mozambique. By leveraging federated estimates, we increase effective sample sizes leading to reduced estimation uncertainty, and enable estimation when single data owners cannot share the data but only have access to the treated or untreated group.

## Introduction

Difference-in-Differences (DID) is a key tool to evaluate treatment effects in fields like finance [Molyneux et al., 2019, Nawaz et al., 2021], clinical research [Galiani et al., 2005, Miller and Wherry, 2017], epidemiology [Goodman-Bacon and Marcus, 2020, Groeniger et al., 2021], and economics [Colchero et al., 2016, Wen et al., 2015]. Recent advances have addressed continuous treatment effects [Callaway et al., 2021], variations in treatment timing [Goodman-Bacon, 2021, Callaway and Sant’Anna, 2021], and varying effects across multiple periods [Callaway and Sant’Anna, 2021]. The Callaway and Sant’Anna [2021] method is available in R [Callaway and Sant’Anna, 2021] and Stata [Rios-Avila et al., 2023], referred to as CSDID (Callaway and Sant’Anna Difference-in-Differences). CSDID has been applied to policy evaluations, such as interventions on teen births [Mark and Wu, 2022] and electric vehicle adoption in response to improved charging infrastructure [Schulz and Rode, 2022], but its use with sensitive individual-level data, like patient data or test performances, has been limited.

In cases involving sensitive individual-level data, sharing is restricted by legal frameworks like the General Data Protection Regulation (GDPR) [Hansen et al., 2021]. This restriction poses 3 major limitations to the analysis: (i) Obtaining consent from all potential participants is challenging yielding reduced sample sizes. (ii) Public health policies often vary regionally, and some entities, like schools or regional governments, may only have data for treated or untreated individuals. If this data cannot be shared, the CSDID cannot be used in these cases. (iii) Students with poorer performance may withhold data due to privacy concerns, introducing selection bias and limiting CSDID’s utility with central learning (Figure 1A). Federated Learning [McMahan et al., 2017] addresses these limitations by enabling multiple data owners to collaborate while iteratively sharing only summary statistics, facilitating joint model training with larger sample sizes and preserving privacy (Figure 1B). It yields parameter estimates with convergence properties identical to central learning, often matching centrally stored data estimates. For CSDID, Federated Learning can overcome the selection bias and increase sample sizes yielding enhanced statistical power. Federated Learning has been successfully applied in various biomedical studies [Dayan et al., 2021, Harrison et al., 2020, Jannasch et al., 2022, Li et al., 2020], but its application to student test performance data has not been explored.

**Figure 1:**
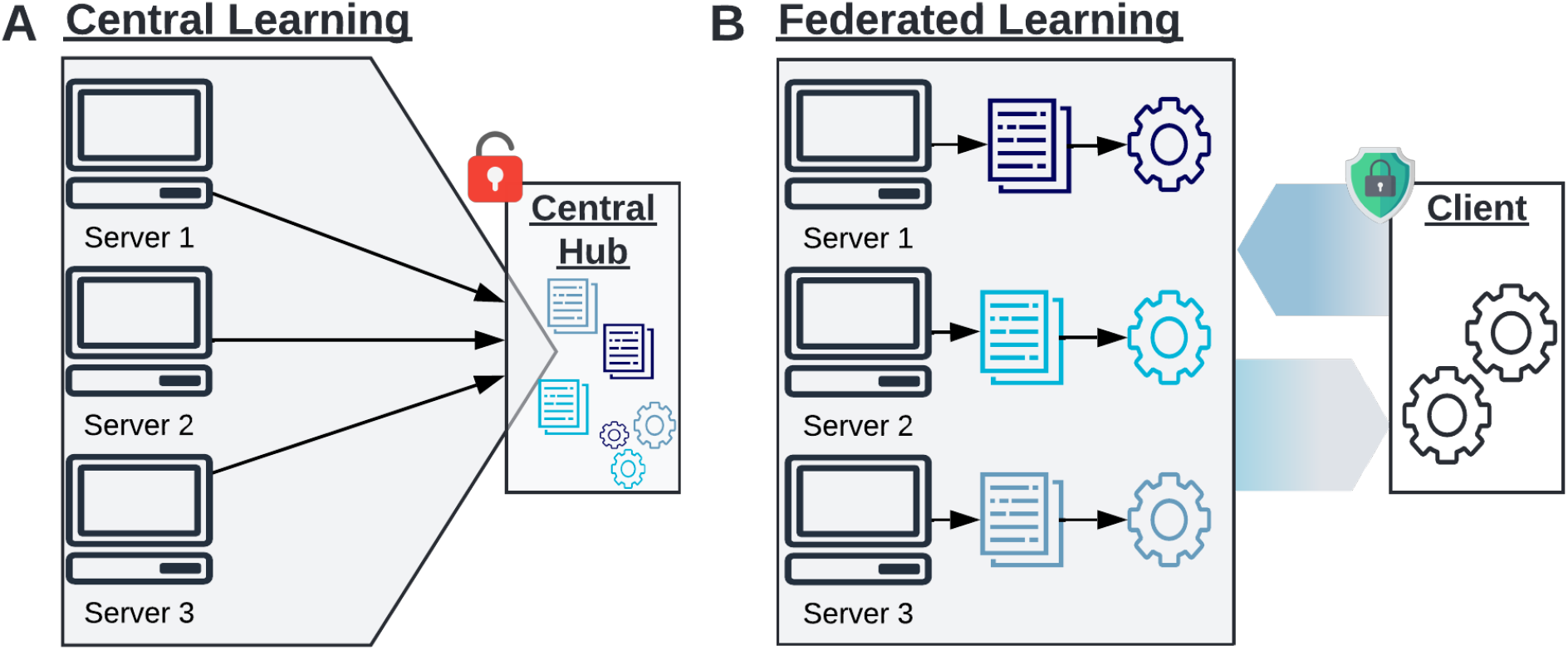
Learning paradigma. **A** Explanation of Central Learning. In Central Learning, models are trained at one central hub (analyst). All servers (data owners) send their data to the central hub, granting the analyst full access to individual-level information without preserving data privacy. **B** Explanation of Federated Learning. In Federated Learning, model updates are computed locally at the servers, and only aggregated information is sent to the analyst (client). The locally aggregated information is then further aggregated to compute overall model parameter updates, which are sent back to the servers. This iterative process continues until a convergence criterion is met, ensuring efficient and privacy-preserving collaboration between servers and the analyst. This figure has been designed using resources from Flaticon.com

We derived a Federated Learning algorithm of the CSDID approach and developed a computational package in DataSHIELD. DataSHIELD [Marcon et al., 2021] is a well-established toolbox for Federated Learning, implemented in the statistical computing language R [R Core Team, 2022]. It is widely used in the biomedical community, including projects like the European Union’s Horizon 2020 ORCHESTRA project and unCover [Tacconelli et al., 2022]. Although DataSHIELD supports many biomedical analysis tools, such as Survival Analysis [Banerjee et al., 2022] and deep Boltzmann Machines [Lenz et al., 2021], a tool for CSDID was previously missing. Our federated tool can obtain exact federated treatment effects, asymptotic standard errors, and distributional equivalent bootstrapped standard errors, comparable to the non-federated implementation [Callaway and Sant’Anna, 2021]. Our software incorporates standard DataSHIELD security measures, such as validity checks on the minimum non-zero counts of observational units and the maximum number of parameters in a regression. Additionally, we have included further security measures tailored to CSDID to prevent attacks as described by Huth et al. [2023]. These measures ensure that computations are secure and confidential, protecting sensitive data from unauthorized access or use.

In the remainder of this paper, we outline the fundamentals of DataSHIELD, explain the federated algorithm, detail the additional security measures implemented in our software, and demonstrate its application through a simulation study. Additionally, we present a case study examining the impact of a malaria elimination initiative on school outcomes, previously analyzed in a non-federated setup [Cirera et al., 2022].

## Results

### DataSHIELD provides federated infrastructure

DataSHIELD [Gaye et al., 2014, Wilson et al., 2017] is an advanced distributed learning ecosystem that utilizes Federated Learning and federated meta-analysis to enable secure and collaborative data analysis across multiple data owners without sharing individual-level data. This approach maintains high statistical power by leveraging larger sample sizes while ensuring data privacy. DataSHIELD’s federated algorithms operate through a client-server structure (Figure 1B): client-side packages run on the analyst’s machine, and server-side packages are installed on the data owner’s machines.

Server-side functions aggregate data locally, detect disclosure risks such as small sample sizes, and send non-disclosive information to the analyst to update parameters or compute overall aggregated statistics.

The dsBase package [Marcon et al., 2022] is the core of the DataSHIELD system, offering a wide range of functionalities, including summary statistics computation and generalized linear models. Additionally, user-written packages extend DataSHIELD’s capabilities, such as restricted Boltzmann machines [Lenz et al., 2021], Omics analysis [Gonzalez et al., 2021], and mediation methods [Avraam and Wheater, 2021]. Our implementation in DataSHIELD enhances this framework by providing a robust tool for evaluating causal impact and treatment effects through the federated version of the CSDID. It is designed to minimize information transfer between the client side and server side, reducing both data leakage risks and communication overhead. Our software package consists of a client-side [Huth, 2023a] and a server-side [Huth, 2023b] R package. The server-side package can be accessed through the client-side package that uses Opal [OBiBa, 2022] for communication.

### Privacy-preserving point estimates are obtained using federated averaging

We present the methodology for computing federated point estimates of the average treatment effect of the treated (ATT). For clarity and consistency, we adopt the notation from Callaway and Sant’Anna [2021].

A dataset suitable for the CSDID analysis comprises multiple evaluation periods *t* and treatment periods *g* (Figure 2A). An ATT and its standard error are computed for each combination of *t* and *g* (Figure 2B) using appropriate control and treatment groups. As the control group, we can use either never-treated or not-yet-treated individuals. While this choice affects the specific individuals used to compute the counterfactual, it does not alter the overall methodology. Therefore, we illustrate the algorithm using never-treated individuals as the control group. Within the package, there are three methods to compute the ATT: the doubly robust (DR) approach, inverse-probability weighting (IPW), and the outcome regression (OR) approach. However, since the IPW and OR approaches are nested within the DR approach, federations are analogous. Thus, detailing the federation of the DR approach suffices.

**Figure 2:**
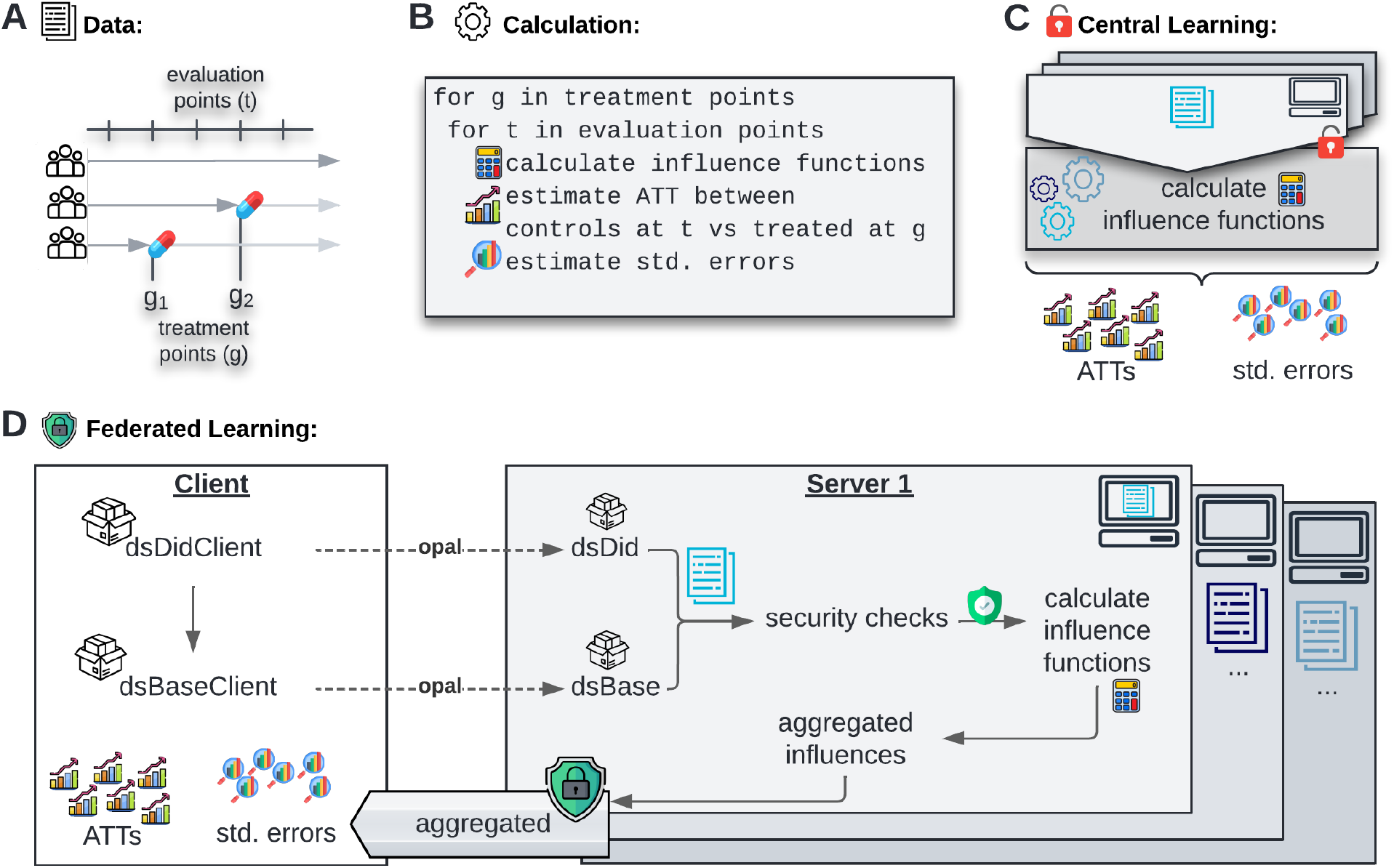
Federated implementation. **A** Visualization of data structure. The data has a panel structure with many observations per individual and varying treatment timing. **B** High-level algorithm of the CSDID. For each combination of evaluation period and treatment period, an ATT and its standard error is computed using the influence function of each individual. **C** CSDID for central learning. In central learning, the data is all at one server such that the sample analogues can be computed directly. **D** CSDID for Federated Learning. The analysis is initalized by using functions from the client side package dsDidClient which calls the dsDid package on the server sides using opal. During the local computations of the influence functions on the server sides, security checks are enforced in order to guarantee data privacy. The server-side influences are aggregated on each server and only the aggregated information is sent to the client side at which ATTs and standard errors are computed. This figure has been designed using resources from Flaticon.com

The non-federated DR ATT is defined as the expectation of the difference in outcome between the treated and the control group, weighted by the probability of receiving treatment. It is given, as described in [Callaway and Sant’Anna, 2021], by

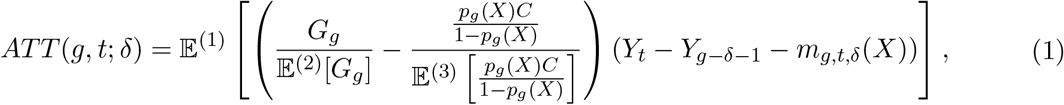

where expectations are donated with a superscript for referencing purposes. Further variables are defined as follows: *δ* is the number of anticipation periods in which individuals know about the upcoming treatment, *G*_*g*_ is a binary variable indicating treatment status in period *g, C* is a binary variable indicating membership in the never-treated control group, *Y*_*t*_ is the outcome variable in period *t*, and *X* is a matrix of pre-treatment covariates. The term *m*_*g*,*t*,*δ*_(*X*) = 𝔼 [*Y*_*t*_ − *Y*_*g*−*δ*−1_|*X, C* = 1] is the expected difference in outcome of the control group and *p*_*g*_(*X*) = 𝔼 [*G*_*g*_|*X, G*_*g*_ + *C* = 1] is the probability of receiving treatment in period *g* given pre-treatment covariates and treatment status.

To estimate the ATT, the expectations in (1) are replaced by their corresponding sample analogues, which we subsequently denote by hat variables. In the case of central learning [Callaway and Sant’Anna, 2021], the data is shared and stored centrally (Figure 2C) allowing all information (*Y*_*t*_, *Y*_*g*−*δ*−1_, *X, G*_*g*_, *C*) to be available for directly computing the sample analogues of all expectations. However, in the federated setting, the data remains on the server sides to avoid sharing sensitive information. Initially, the servers (but not the client) have access to their individual data (*Y*_*t*_, *Y*_*g*−*δ*−1_, *X, G*_*g*_, *C*). Thus, the sample analogues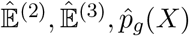, and 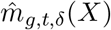 can be computed by the client using federated means and federated generalized linear models (GLMs) (Figure 2D) via dsBase. For the federated means, only server-aggregated means need to be shared and for the federated GLMs iterative gradient sharing is used to obtain GLM parameter estimates. These parameters and means are then sent to the servers to compute the sample analogue of the term inside the expectation of 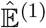 locally. These local influences can be aggregated using a federated mean in order to compute 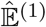 on the client-side. A more detailed description of the computation of the federated sample analogues, federated influence functions as well as a description of the federated standard error algorithms is given in the Method Details section.

### Additional privacy-enhancing measures increase security

Our package adheres to DataSHIELD’s data security standards and the highest security standards as per recent assessments, ensuring all function outputs comply with the disclosure settings configured in Opal [OBiBa, 2022]. These measures include, inter alia, preventing the subsetting of data frames if the subset does not meet the minimum row requirement and prohibiting aggregations with fewer objects than the specified threshold. Given that CSDID subsets individuals for each treatment period and a fixed evaluation period (Figure 2B), servers with low sample sizes may not meet DataSHIELD’s security requirements and, therefore, cannot participate in computing the ATT for those periods. To address this, our package automatically checks in advance which servers have sufficient observations for the respective periods and excludes only those servers for the respective computations. This ensures that treatment effect estimations use larger sample sizes from the remaining servers, improving accuracy and efficiency while maintaining data security and privacy.

Additionally, when performing federated estimation of the influence functions, it is necessary to append columns to a matrix (`ds.AppendInfluence`). However, this process enables malicious clients to append arbitrary columns to a matrix and therefore the creation of known linearly independent matrices which can cause data leakage of all data points [Huth et al., 2023]. To mitigate this risk, we implement a strategy of row-wise shuffling of data frames and matrices after appending columns. This maintains the integrity of results through ID column matching while protecting against data leakage attacks.

Using globally summarized quantities, such as global means, on the server sides requires data transmission from the client. However, unrestricted data transmission can facilitate data leakage attacks. As a countermeasure, we process all data sent to the server immediately, ensuring it is not stored. For data that must be stored, the client is limited to sending a single number at a time to prevent attacks. Sending known numbers to the servers raises the potential threat of data leakage if the client is able to concatenate multiple known single numbers into different linearly independent vectors [Huth et al., 2023]. Therefore, it is not recommended to use this package with `ds.cbind` and `ds.rbind` from DataSHIELD’s base package [Marcon et al., 2022], which can be disabled in Opal to enhance data privacy.

The Method Details section provides further details on the specific functions and the security measures implemented to protect the data privacy.

### Federated estimates preserve privacy and equal central learning estimates

To validate our package, we used a simulated dataset generated with the non-federated R package and compared our federated implementation to the central learning. Specifically, we compared point estimates of the ATT, asymptotic standard errors, and bootstrapped standard errors. Additionally, we assessed the reduction of uncertainty achieved by the federated model compared to study-level analysis, where each data owner analyzes only their local dataset.

We simulated a dataset of 801 individuals with observations at four time periods, randomly allocated across six servers (Figure 3). Three servers contained 134 individuals (536 observations each), and three contained 133 individuals (532 observations each). Individuals were either never treated or treated in period two or three. All observations for an individual were stored on one server, reflecting the realistic scenario where different data providers, such as hospitals or schools, do not share data on the same individual. Our main results used the DR estimator with not-yet-treated individuals as the control group. Qualitatively equivalent results for the IPW and OR methods and other control groups are presented in the Supplementary Material.

**Figure 3:**
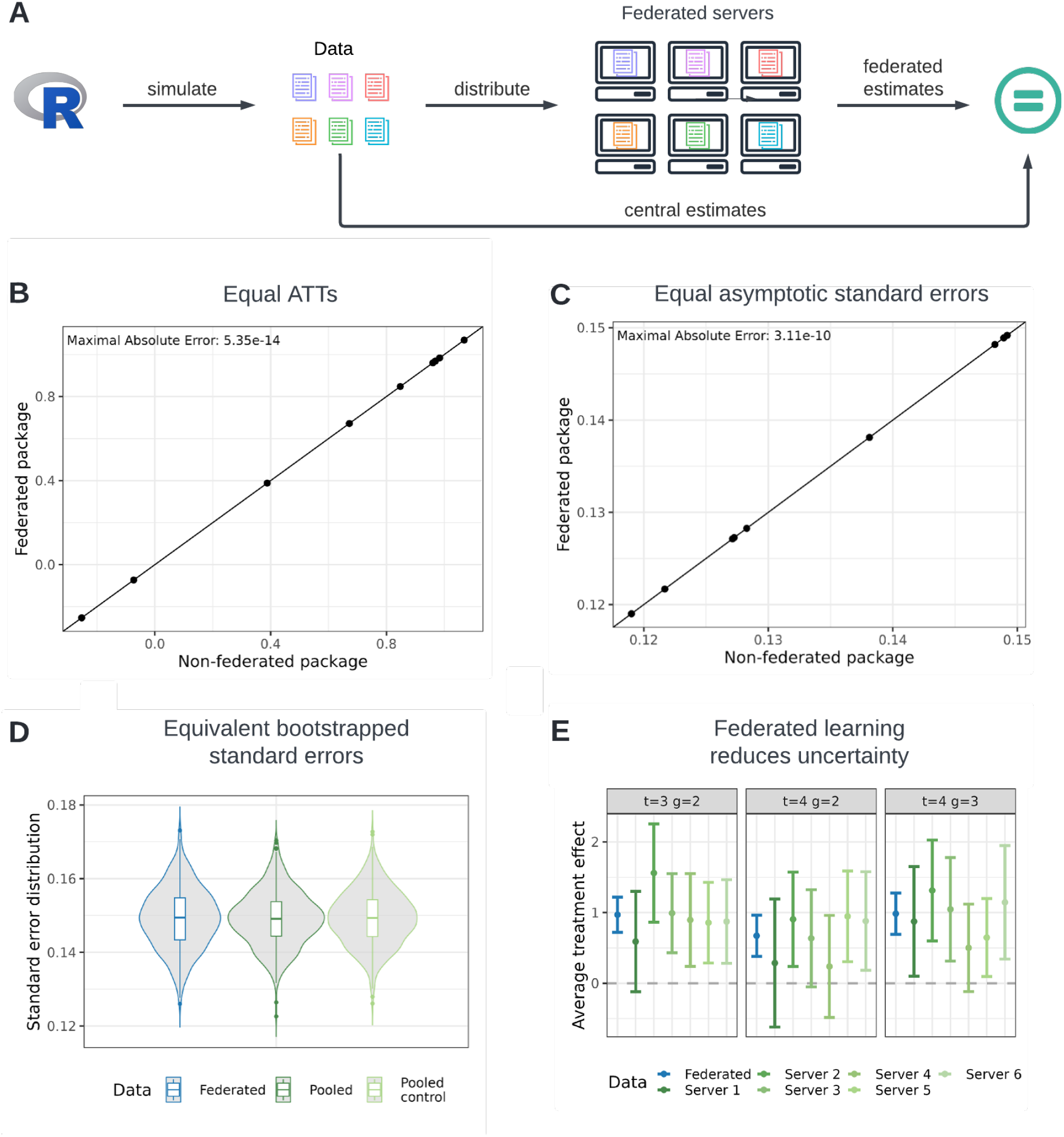
Similarity of Federated and Central Estimation for the DR estimate with not-yet-treated individuals as the control group. **A** Simulation setup. The federated setup involved 6 servers. Three servers had 134 (536) individuals (observations), and three had 133 (532) individuals (observations). Individuals were either never treated or treated in period two or three, with all observations of an individual on one server. **B** Equality of central and federated point estimates. The x-axis shows central (non-federated) estimates [Callaway and Sant’Anna, 2021], while the y-axis shows federated estimates. The 45° line indicates equal results when aligned. **C** Equality of central and federated asymptotic standard errors. The x-axis shows central asymptotic standard errors, and the y-axis shows federated standard errors. The 45° line is shown for reference. **D** Comparison of bootstrapped standard errors. Boxplots and densities compare the distribution of federated (blue) and central (green) bootstrapped standard errors. Two central learning distributions establish reference differences, with 500 bootstrapped standard errors analyzed. **E** Treatment effect estimates. Point estimates (dots) and 95% confidence intervals for post-treatment periods are shown. Federated package estimates are in blue; estimates from one server are in green. This figure has been designed using resources from Flaticon.com

The federated package successfully reproduced the average treatment effect (Figure 3B) and asymptotic standard errors (Figure 3C) up to numerical precision. For the federated bootstrapped standard errors, we relied on visual inspection to evaluate empirical validity, as statistical tests could only reject the hypothesis of equal distributions, not that they are unequal. Detailed reasoning for this limitation is provided in the Method Details. We used a Monte Carlo simulation, repeating the bootstrap computation 500 times for both federated and central settings. Empirical distributions were generated, and the difference between two central learning distributions served as a benchmark for comparison. Violin plots and boxplots (Figure 3D) indicated that the quantiles of all distributions are visually almost identical, and the densities show the same shapes, providing strong visual evidence that the federated and central learning bootstrapped standard errors are distributional equivalent.

We also examined the impact of the federated implementation on the estimation of the overall treatment effect compared to study-level analysis. By observing the point estimates and 95% confidence intervals, with standard errors obtained from the multiplier bootstrap (Figure 3E), we found: (i) study-level point estimates fluctuated around the federated and central learning point estimate, and (ii) increased uncertainty in the study-level DR estimates was reflected in wider confidence intervals. The study-level point estimates fluctuated around the federated estimate since the federated simulated data is independent and identically distributed (*i*.*i*.*d*.) in individuals and we allocated the simulated individuals randomly to the servers. Therefore, the server data is also *i*.*i*.*d*, and thus, the point estimates of the study-level analysis fluctuate around the federated estimate. This *i*.*i*.*d*. case is the most favorable scenario for study-level analysis; point estimates will deviate more from the federated estimate in non-*i*.*i*.*d*. settings. The increased size of the confidence intervals reflects reduced sample sizes, resulting in more uncertainty and potentially failing to reject the null hypothesis of no treatment effect in study-level cases where it could be rejected in the federated case. The federated setup yields estimates with lower uncertainty, enhancing the detection of true treatment effects.

### Federated learning enables CSDID analysis in cases when central learning fails

To test the effectiveness of our federated framework in a real-world scenario, we studied the impact of malaria interventions on school outcomes in Mozambique. In 2015, a malaria elimination initiative was implemented in Magude (Figure 4A, B) as a quasi-experiment to estimate its impact on selected primary school outcomes. Data was collected from nine schools in two districts of Mozambique. This data was initially analyzed using a regression DID approach [Cirera et al., 2022], which required data sharing agreements for data transfer to analysts. For our analysis, we conducted a complete case analysis with the CSDID, including data from 1,044 individuals with observations available for the three school terms of 2015 and 2016 (Figure 4B). Sample statistics for each school are provided in Table 2 in Method Details.

**Table 1:** New DataSHIELD functions. implemented in our software package. Functions starting with *ds*. are client-side functions and can be called via the dsDidClient [Huth, 2023a] package.

| Client side function | Description |
| --- | --- |
| `ds.did` | Calculates the average treatment effect of the treated, standard errors, and pre-tests for the parallel trend assumption. |
| `ds.addColumnOnes` | Adds a column of ones to a dataframe on the server side. |
| `ds.appendInfluence` | Appends influence to a specified column for given ids and periods in a dataframe on the server side. |
| `ds.computeMatrixCrossproduct` | Computes cross product of a matrix on the server side. |
| `ds.computeOdds` | Computes odds using propensity scores for a given dataset and outcome variable. |
| `ds.createEmptyIdMatrix` | Creates empty matrix with unique identifiers from a dataframe on the server side. |
| `ds.enoughIndividuals` | Checks if enough individuals in a dataframe have a specified value in a specified column on the server side. |
| `ds.generateNotYetTreated` | Generates a dataframe of untreated individuals based on given variables on the server side. |
| `ds.multiplierBootstrap` | Performs multiplier bootstrap on a given influence matrix on the server side. |
| `ds.multiplyMatrixMatrix` | Multiplies two matrices and assigns result to a new object on the server side. |
| `ds.multiplyMatrixScalar` | Multiplies a matrix and a scalar and assigns result to a new object on the server side. |
| `ds.recode` | Creates a new object and replaces entries of a vector. |
| `ds.sendToServer` | Sends a single number to the server side. |
| `ds.subsetDf` | Subsets a dataframe based on a specified variable and value. |

**Table 2:**
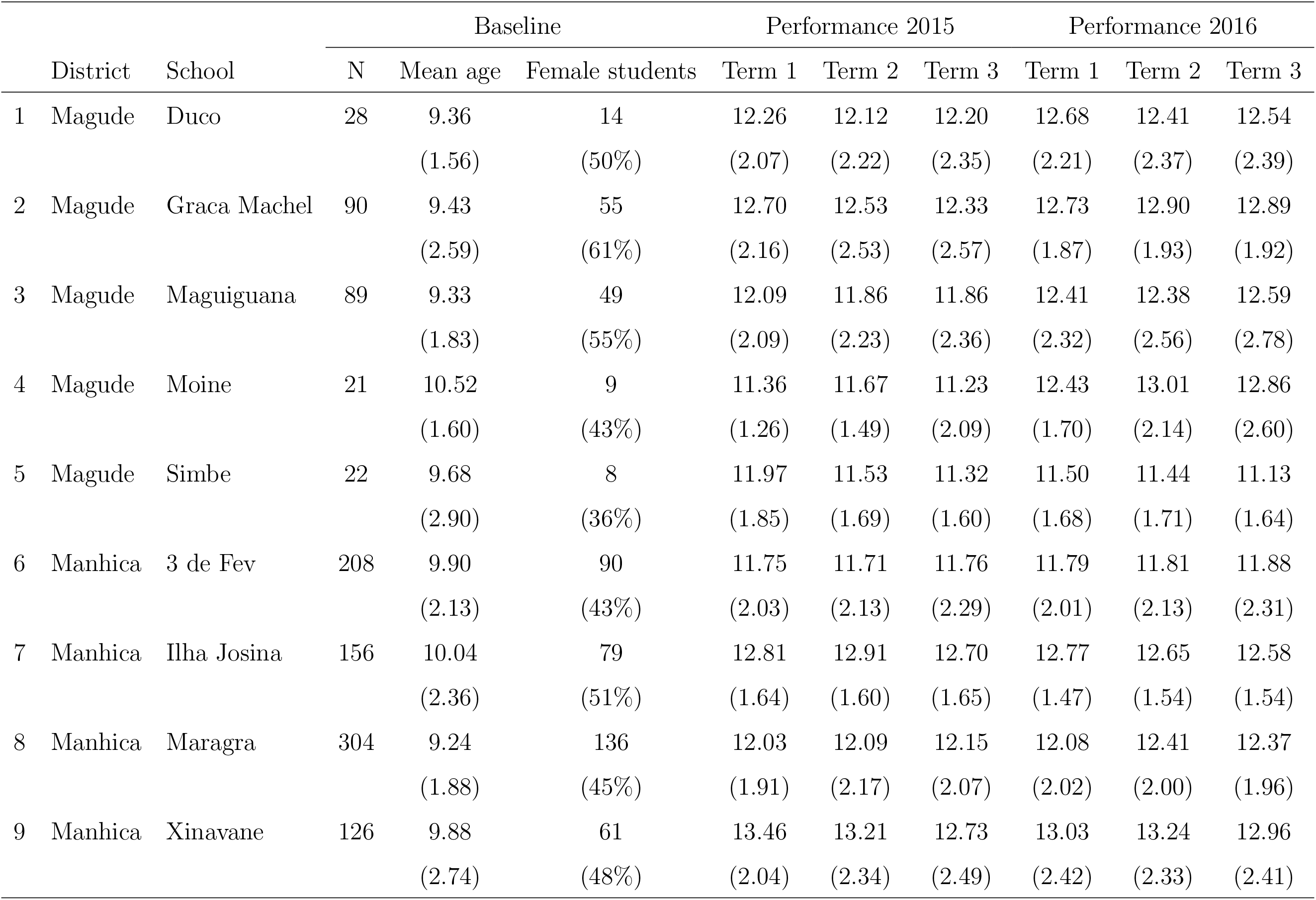
Summary statistics: Average performance of students by district, school and term of studies. Standard deviations are shown in parenthesis for all sample averages. For number of females, the table reports the percentage of female students on the total school population N.

**Figure 4:**
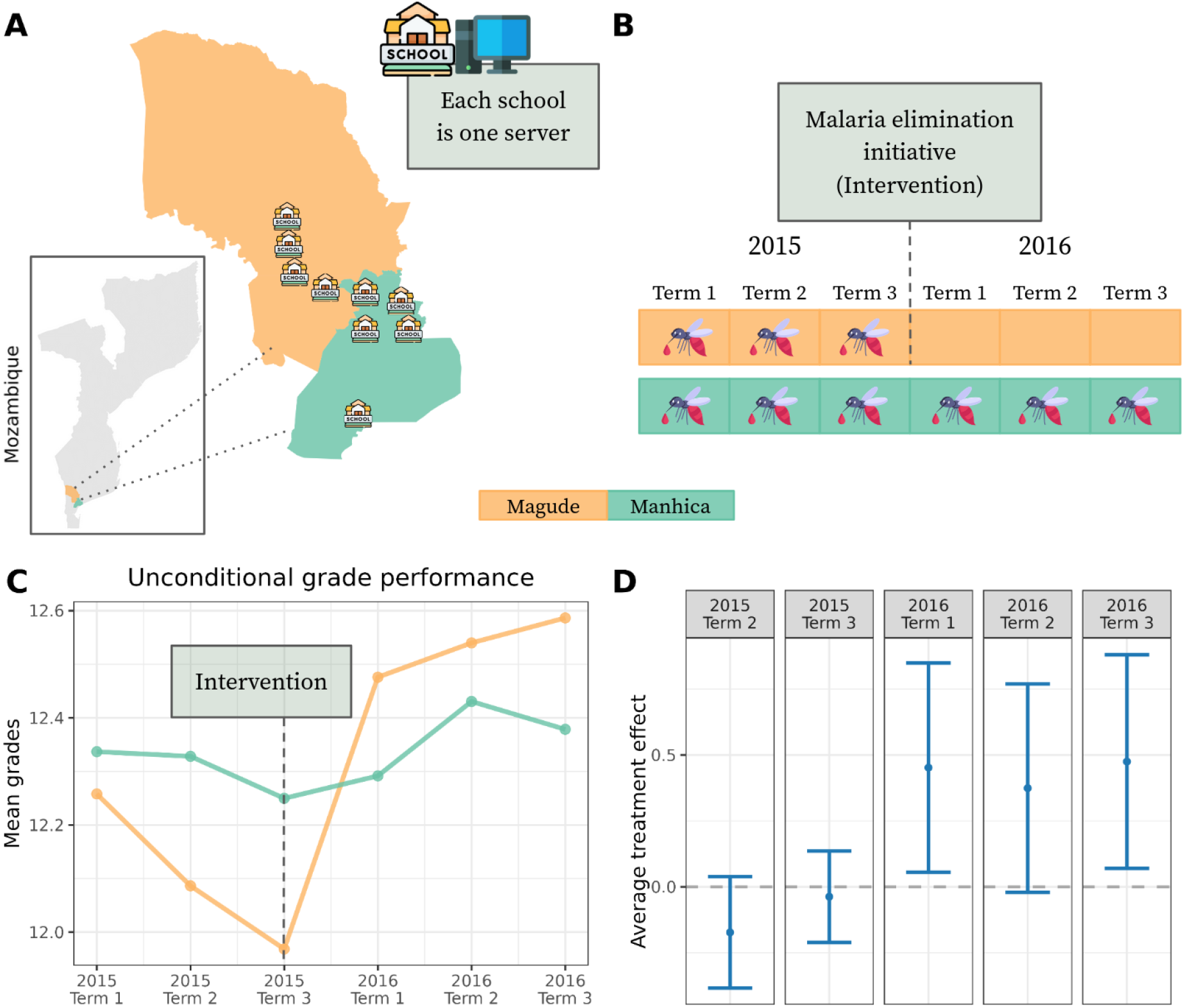
Federated learning enables estimation of DR estimates. **A** The districts of Magude and Manhiça in Mozambique and the location of the 9 schools. **B** The timeline of the malaria elimination initiative that was launched in Magude before the term 1 in 2016. **C** Unconditional means of the mean grades in Magude and Manhiça respectively. The means are estimated as sample averages over the observations (mean grades of students) of the individuals in the respective districts at the respective time. **D** The subplot presents point estimates (depicted as dots) and 95% confidence intervals of the estimated federated treatment effects for all periods. This figure has been designed using resources from Flaticon.com

**Figure 5:**
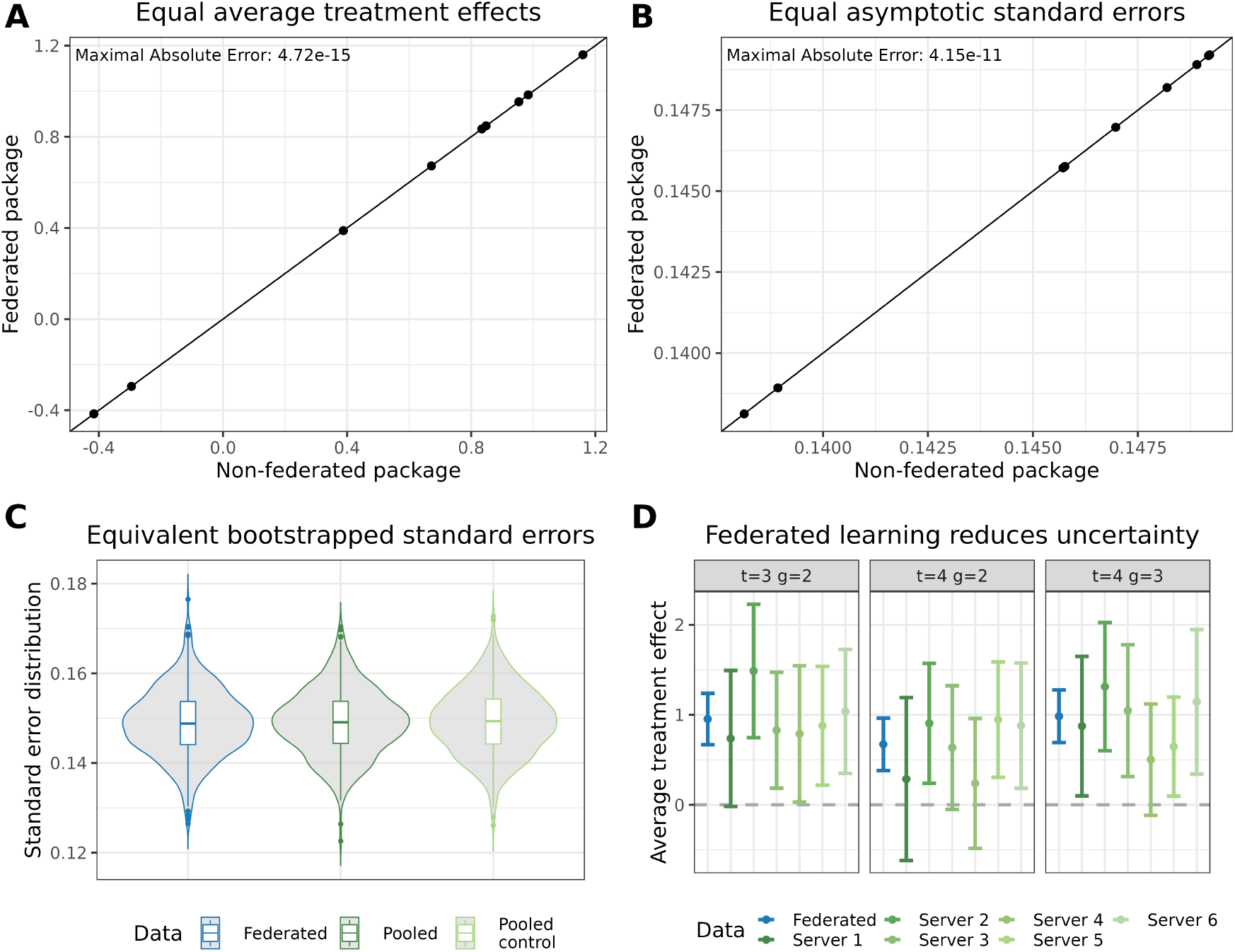
DR estimate and never treated individuals as control group. The federated set-up consisted of 6 servers. Three servers contained 134 (536) individuals (observations), and three servers contained 133 (532) individuals (observations). Individuals were either never treated or treated in period two or three. All observations of one individual were within one server. **A** Depicts the equality of central and federated point estimates. The x-axis represents the point estimates of the central (non-federated) estimator [Callaway and Sant’Anna, 2021], while the y-axis represents the estimates obtained from our federated approach. The diagonal line depicts the 45° line, indicating that the federated estimate yields results equivalent to the non-federated estimate when they align along this line. **B** Depicts the equality of central and federated asymptotic standard errors. The x-axis represents the asymptotic standard errors of the central (non-federated) estimator, while the y-axis represents the asymptotic standard errors obtained from our federated approach. The diagonal line again depicts the 45° line. **C** The subplot displays a comparison of the distribution of bootstrapped standard errors using boxplots and densities. The distribution of the federated bootstrapped standard errors is shown in blue, while the distribution of the central estimation is shown in green (dark and light). Two central learning distributions were computed to establish a plausible reference difference between two equal distributions. A total of 500 bootstrapped standard errors were computed to obtain the distributions for analysis. **D** The subplot presents point estimates (depicted as dots) and 95% confidence intervals of the estimated treatment effects for the periods following the treatment. The estimates obtained with our federated package are shown in blue, while estimates based on data from only one server are depicted in green. This figure has been designed using resources from Flaticon.com

**Figure 6:**
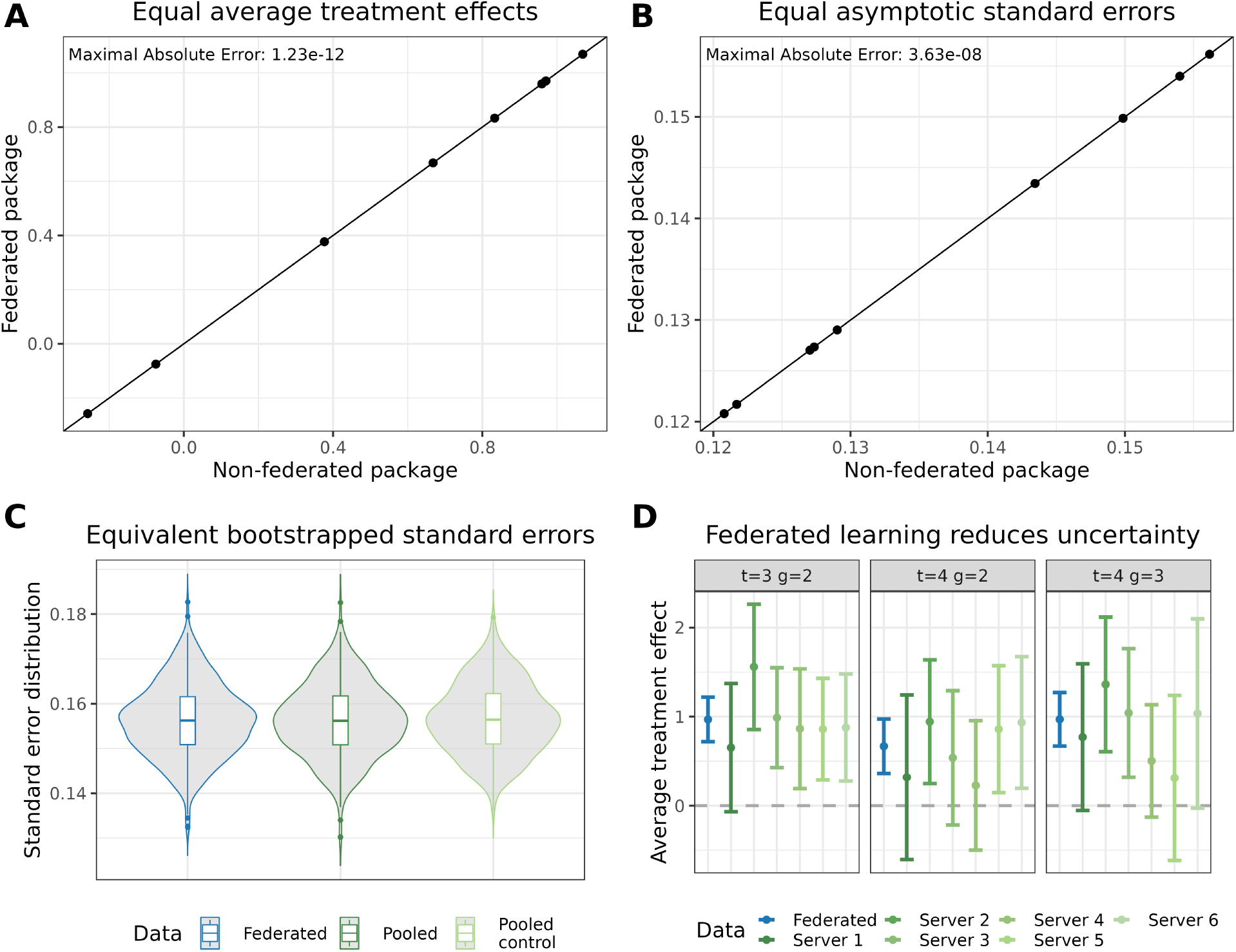
Inverse probability weighted estimate and not-yet treated individuals as control group. The federated set-up consisted of 6 servers. Three servers contained 134 (536) individuals (observations), and three servers contained 133 (532) individuals (observations). Individuals were either never treated or treated in period two or three. All observations of one individual were within one server. **A** Depicts the equality of central and federated point estimates. The x-axis represents the point estimates of the central (non-federated) estimator [Callaway and Sant’Anna, 2021], while the y-axis represents the estimates obtained from our federated approach. The diagonal line depicts the 45° line, indicating that the federated estimate yields results equivalent to the non-federated estimate when they align along this line. **B** Depicts the equality of central and federated asymptotic standard errors. The x-axis represents the asymptotic standard errors of the central (non-federated) estimator, while the y-axis represents the asymptotic standard errors obtained from our federated approach. The diagonal line again depicts the 45° line. **C** The subplot displays a comparison of the distribution of bootstrapped standard errors using boxplots and densities. The distribution of the federated bootstrapped standard errors is shown in blue, while the distribution of the central estimation is shown in green (dark and light). Two central learning distributions were computed to establish a plausible reference difference between two equal distributions. A total of 500 bootstrapped standard errors were computed to obtain the distributions for analysis. **D** The subplot presents point estimates (depicted as dots) and 95% confidence intervals of the estimated treatment effects for the periods following the treatment. The estimates obtained with our federated package are shown in blue, while estimates based on data from only one server are depicted in green. This figure has been designed using resources from Flaticon.com

**Figure 7:**
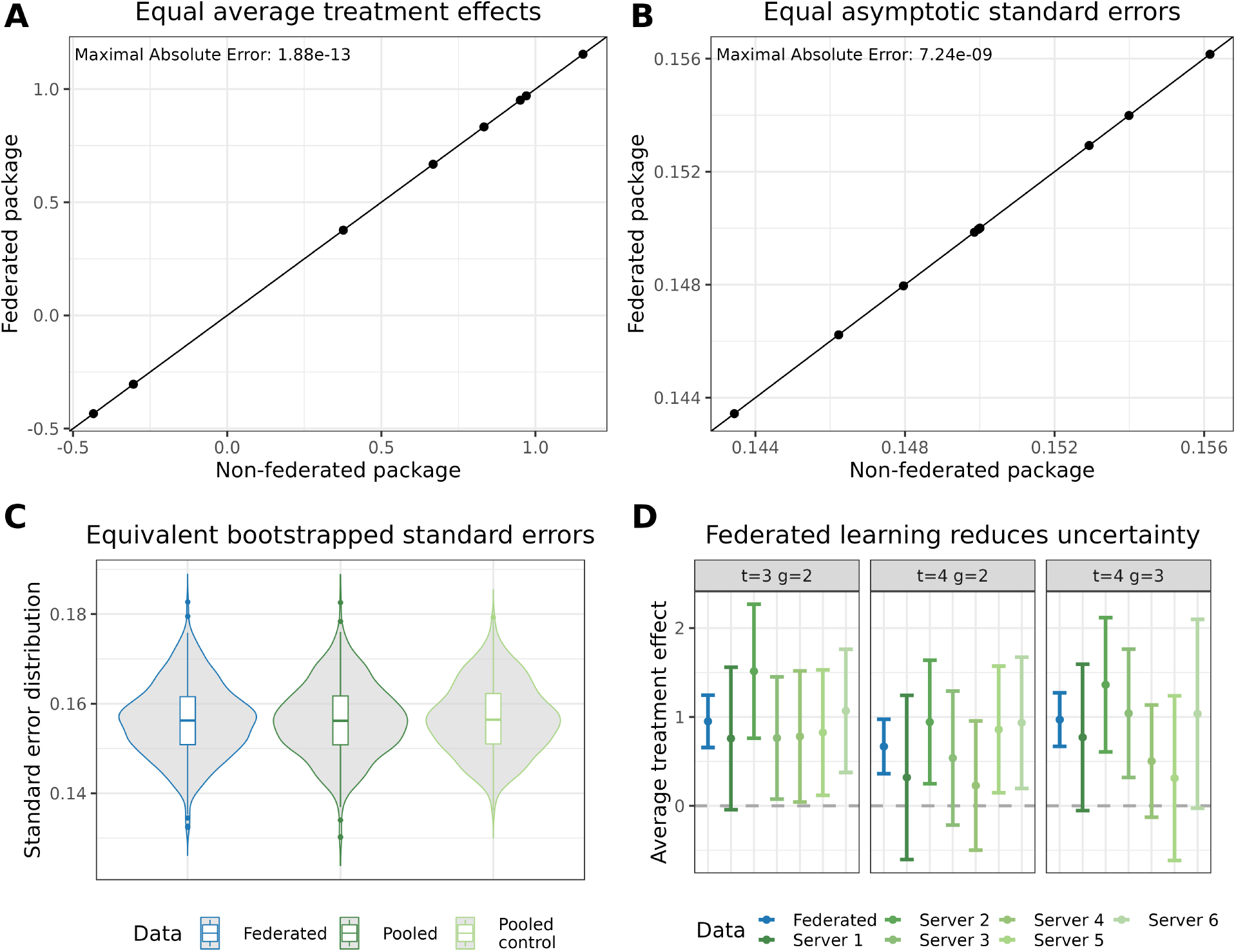
Inverse probability weighted estimate and never treated individuals as control group. The federated set-up consisted of 6 servers. Three servers contained 134 (536) individuals (observations), and three servers contained 133 (532) individuals (observations). Individuals were either never treated or treated in period two or three. All observations of one individual were within one server. **A** Depicts the equality of central and federated point estimates. The x-axis represents the point estimates of the central (non-federated) estimator [Callaway and Sant’Anna, 2021], while the y-axis represents the estimates obtained from our federated approach. The diagonal line depicts the 45° line, indicating that the federated estimate yields results equivalent to the non-federated estimate when they align along this line. **B** Depicts the equality of central and federated asymptotic standard errors. The x-axis represents the asymptotic standard errors of the central (non-federated) estimator, while the y-axis represents the asymptotic standard errors obtained from our federated approach. The diagonal line again depicts the 45° line. **C** The subplot displays a comparison of the distribution of bootstrapped standard errors using boxplots and densities. The distribution of the federated bootstrapped standard errors is shown in blue, while the distribution of the central estimation is shown in green (dark and light). Two central learning distributions were computed to establish a plausible reference difference between two equal distributions. A total of 500 bootstrapped standard errors were computed to obtain the distributions for analysis. **D** The subplot presents point estimates (depicted as dots) and 95% confidence intervals of the estimated treatment effects for the periods following the treatment. The estimates obtained with our federated package are shown in blue, while estimates based on data from only one server are depicted in green. This figure has been designed using resources from Flaticon.com

**Figure 8:**
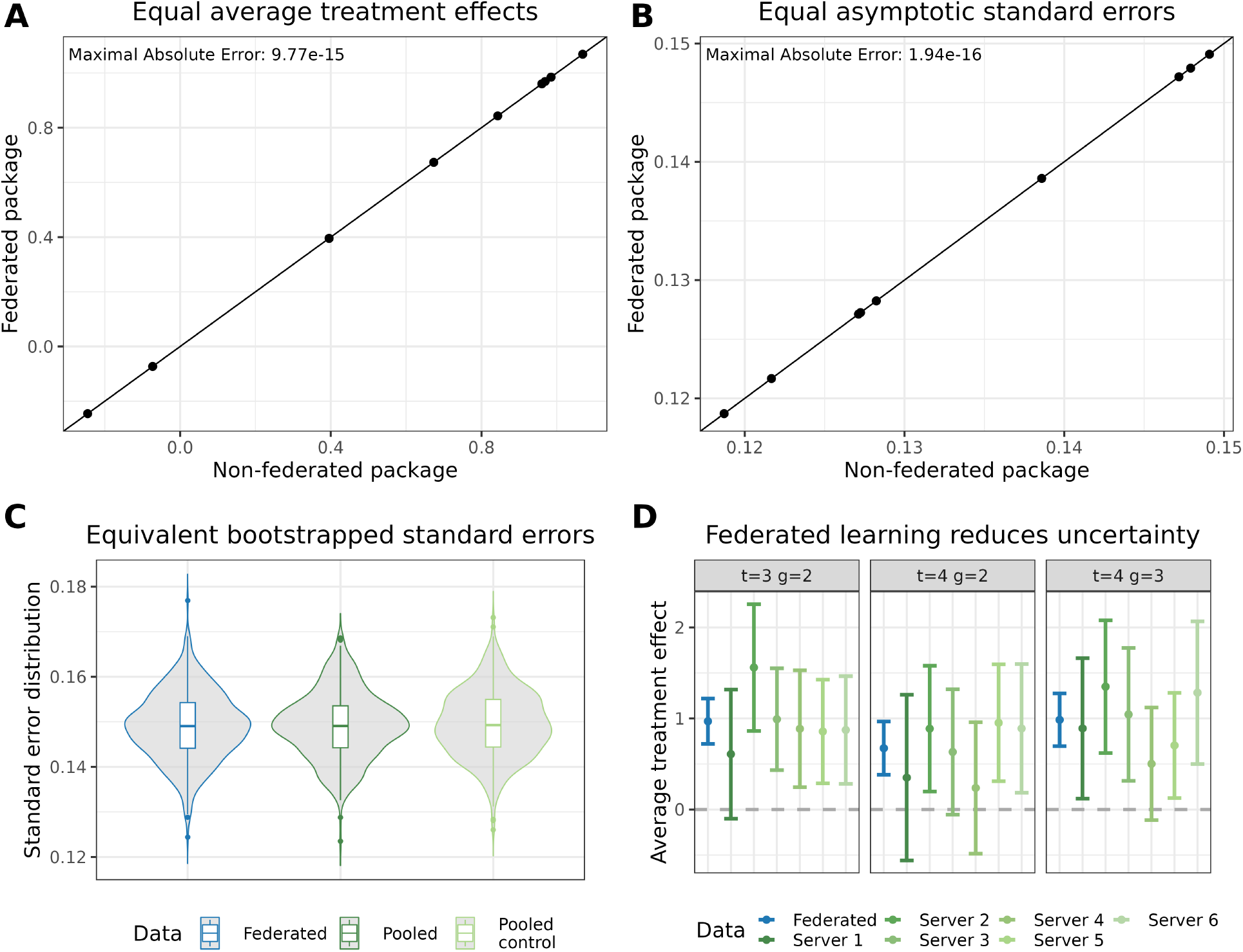
Outcome regression estimate and not-yet treated individuals as control group. The federated set-up consisted of 6 servers. Three servers contained 134 (536) individuals (observations), and three servers contained 133 (532) individuals (observations). Individuals were either never treated or treated in period two or three. All observations of one individual were within one server. **A** Depicts the equality of central and federated point estimates. The x-axis represents the point estimates of the central (non-federated) estimator [Callaway and Sant’Anna, 2021], while the y-axis represents the estimates obtained from our federated approach. The diagonal line depicts the 45° line, indicating that the federated estimate yields results equivalent to the non-federated estimate when they align along this line. **B** Depicts the equality of central and federated asymptotic standard errors. The x-axis represents the asymptotic standard errors of the central (non-federated) estimator, while the y-axis represents the asymptotic standard errors obtained from our federated approach. The diagonal line again depicts the 45° line. **C** The subplot displays a comparison of the distribution of bootstrapped standard errors using boxplots and densities. The distribution of the federated bootstrapped standard errors is shown in blue, while the distribution of the central estimation is shown in green (dark and light). Two central learning distributions were computed to establish a plausible reference difference between two equal distributions. A total of 500 bootstrapped standard errors were computed to obtain the distributions for analysis. **D** The subplot presents point estimates (depicted as dots) and 95% confidence intervals of the estimated treatment effects for the periods following the treatment. The estimates obtained with our federated package are shown in blue, while estimates based on data from only one server are depicted in green. This figure has been designed using resources from Flaticon.com

**Figure 9:**
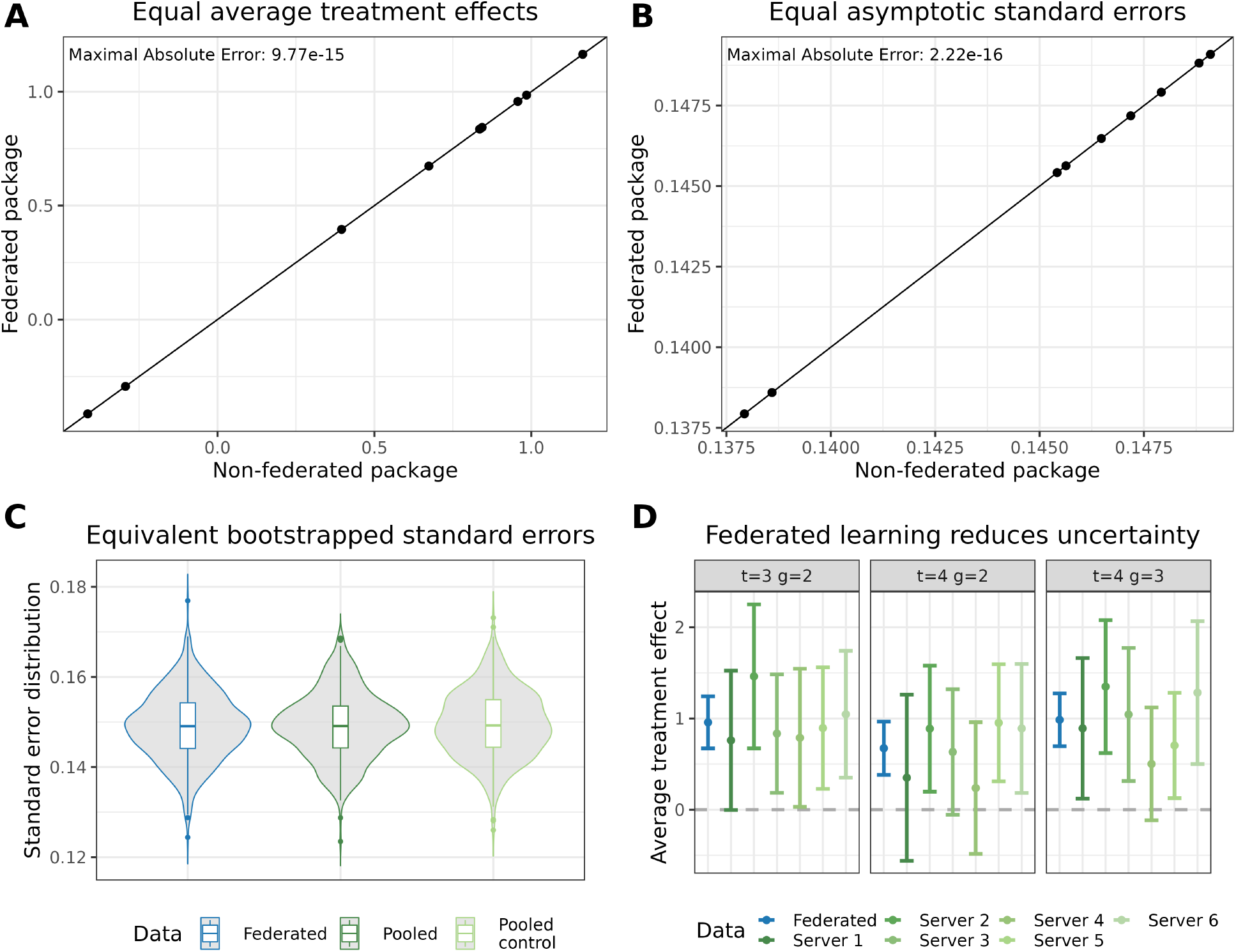
Outcome regression estimate and never treated individuals as control group. The federated set-up consisted of 6 servers. Three servers contained 134 (536) individuals (observations), and three servers contained 133 (532) individuals (observations). Individuals were either never treated or treated in period two or three. All observations of one individual were within one server. **A** Depicts the equality of central and federated point estimates. The x-axis represents the point estimates of the central (non-federated) estimator [Callaway and Sant’Anna, 2021], while the y-axis represents the estimates obtained from our federated approach. The diagonal line depicts the 45° line, indicating that the federated estimate yields results equivalent to the non-federated estimate when they align along this line. **B** Depicts the equality of central and federated asymptotic standard errors. The x-axis represents the asymptotic standard errors of the central (non-federated) estimator, while the y-axis represents the asymptotic standard errors obtained from our federated approach. The diagonal line again depicts the 45° line. **C** The subplot displays a comparison of the distribution of bootstrapped standard errors using boxplots and densities. The distribution of the federated bootstrapped standard errors is shown in blue, while the distribution of the central estimation is shown in green (dark and light). Two central learning distributions were computed to establish a plausible reference difference between two equal distributions. A total of 500 bootstrapped standard errors were computed to obtain the distributions for analysis. **D** The subplot presents point estimates (depicted as dots) and 95% confidence intervals of the estimated treatment effects for the periods following the treatment. The estimates obtained with our federated package are shown in blue, while estimates based on data from only one server are depicted in green. This figure has been designed using resources from Flaticon.com

In our federated setup, each of the nine schools hosted their own server, simulating that student data never left the school premises (Figure 4A). Consequently, the schools in Magude (four schools) only had data on treated individuals from 2015, while the schools in Manhiça (five schools) only had data on control group individuals. This separation would make it impossible to estimate non-federated local CSDID average treatment effects without sharing the data with a central hub.

Inspecting the unconditional means, computed in a federated manner, we observed that before the malaria elimination initiative, Magude (the treated district) had a lower grade average compared to Manhiça (Figure 4C). After the initiative, student performance in Magude improved significantly, whereas the improvement in Manhiça was comparatively moderate. To test the hypothesis of the malaria intervention’s effect, we ran the federated CSDID with age and gender as covariates to control for potential confounders (Figure 4D). The results revealed significant differences (at the 95% confidence level) in the first and third terms of 2016, while the estimate for the second term was close to the boundary of significance.

## Discussions

The use of Federated Learning has emerged as a promising approach to increase sample sizes when dealing with sensitive data. However, the availability of tools for causal analysis within this context has been limited. To address this gap and make such methods accessible to the research community, we developed a federated version of the Difference-in-Differences estimator proposed by Callaway and Sant’Anna [2021] and implemented it in the Federated Learning platform DataSHIELD us-ing the statistical programming language R. Our federated package covers many functionalities of the CSDID R package [Callaway and Sant’Anna, 2021] and additionally allows for the estimation of treatment effects when treatment and control groups are only available mutually exclusively to single data owners. Our results demonstrate that the proposed federated version of the CSDID estimator can be implemented while preserving the original estimates, asymptotic standard errors, and distributional equivalent bootstrapped standard errors. Additionally, Federated Learning reduces uncertainty and allows estimation in scenarios where treated and untreated individuals are separated across data owners. We have ensured that standard DataSHIELD security measures are in place to protect data privacy and have further enhanced security with additional measures.

We specifically chose the Callaway and Sant’Anna estimator due to its high relevance and flexibility. This estimator accommodates multiple time periods for treatments and evaluations, incorporates different types of control groups, and offers various estimators and simultaneous confidence bands via bootstrapped standard errors. Moreover, it requires the parallel trend assumption to hold only after conditioning on covariates, relaxing the assumption of the classical regression DID where the parallel trend must hold unconditionally. Future work on federated causal impact analysis could explore estimating treatment effects from repeated cross-sections in the context of CSDID, as implemented in Callaway and Sant’Anna [2021], or synthetic control methods that have gained traction in recent years [Abadie et al., 2010, 2015].

Our federated version of the CSDID fills a gap in the availability of tools for causal analysis in Federated Learning. Our results highlight the potential of using Federated Learning for privacy-preserving causal analysis in settings with sensitive data, while maintaining high levels of data security through standard and additional measures implemented within DataSHIELD.

### Limitations of the study

This study has two main limitations. The first significant limitation, shared with the non-federated method, is the necessity to bin the time points of individual records into discrete events to apply the CSDID. This binning process can result in the loss of precise information regarding treatment timings, especially when dealing with data from non-equidistant observations. However, the federated approach allows for the incorporation of more data points, which can potentially lead to a finer granularity in event discretization. Secondly, the use of this software package requires the establishment of a client-server infrastructure. This setup can be challenging for users without extensive technical experience. Nevertheless, the DataSHIELD platform provides comprehensive tutorials and community support, which can assist users in overcoming these technical hurdles and adapting to the process more efficiently.

## Abbreviations

ATT: Average treatment effect of the treated
CSDID: Callaway and Sant’Anna Difference-in-Differences
DID: Difference-in-Differences
DR: Doubly Robust
IPW: Inverse probability weighting
RO: Regression Outcome
SE: Standard error

## Acknowledgements

This study was funded by the German Research Foundation (Deutsche Forschungsgemeinschaft, DFG) under Germany’s Excellence Strategy (EXC 2047 - 390685813 & EXC 2151 - 390873048) and under the project IDs 432325352 – SFB 1454 and 458597554 – SEPAN, the University of Bonn (via the Schlegel Professorship of JH), the Helmholtz Association - Munich School for Data Science (MUDS), and the ORCHESTRA project. The ORCHESTRA project has received funding from the European Union’s Horizon 2020 research and innovation program under grant agreement No 101016167. The views expressed in this paper are the sole responsibility of the authors and the Commission is not responsible for any use that may be made of the information it contains. The funders had no role in the study design, data collection, data analyses, data interpretation, writing, or submission of this manuscript.

The icons for the figures were created by Freepik, Flat Icons, Roundicons, Smashicons, and smash-ingstocks from www.flaticon.com.

## Authors contributions

M.H. developed the federation of the algorithms. M.H. and C.G, implemented the software packages in R. L.C., E.S., F.S. collected the data for the study in Mozambique. M.H. and L.S. visualized the results. J.H and E.S. conceptualized the study. M.H., E.S. and J.H. wrote the manuscript. All authors read and approved the final manuscript.

## Declaration of generative AI and AI-assisted technologies in the writing process

During the preparation of this work the authors used ChatGPT4 in order to improve the readability and language of the manuscript. After using ChatGPT4, the authors reviewed and edited the content as needed and take full responsibility for the content of the published article.

## Star methods

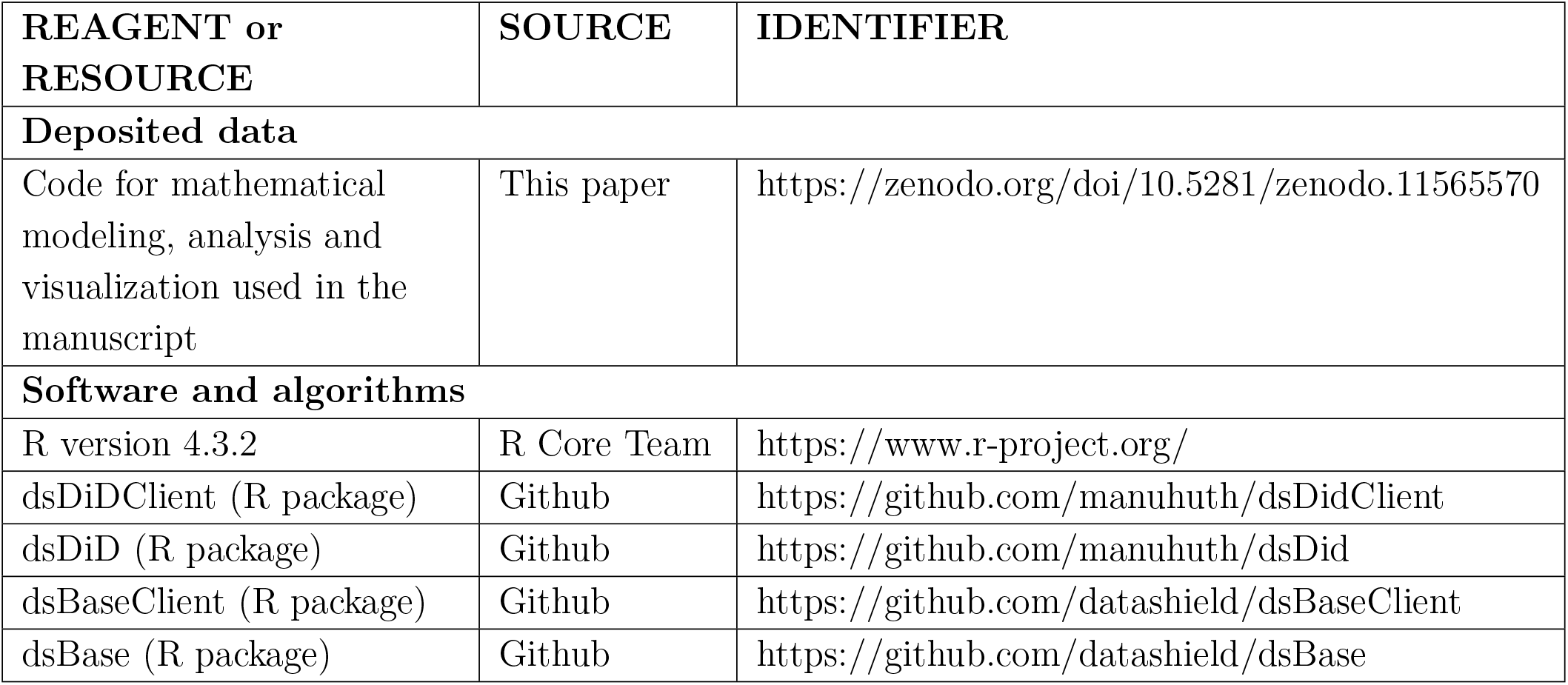

| REAGENT or RESOURCE | SOURCE | IDENTIFIER |
| --- | --- | --- |
| **Deposited data** |  |  |
| Code for mathematical modeling, analysis and visualization used in the manuscript | This paper | https://zenodo.org/doi/10.5281/zenodo.11565570 |
| **Software and algorithms** |  |  |
| R version 4.3.2 | R Core Team | https://www.r-project.org/ |
| dsDiDClient (R package) | Github | https://github.com/manuhuth/dsDidClient |
| dsDiD (R package) | Github | https://github.com/manuhuth/dsDid |
| dsBaseClient (R package) | Github | https://github.com/datashield/dsBaseClient |
| dsBase (R package) | Github | https://github.com/datashield/dsBase |

## Resource availability

### Lead contact

Further information and requests for resources should be directed to and will be fulfilled by the lead contact, Jan Hasenauer

### Materials availability

This study did not generate new unique reagents.

### Data and code availability

- This paper analyses data in the example of the CSDID. Use of the Mozambique school data requires a data sharing agreement between your institution and the Centro de Investigação em Saúde de Manhiça as it contains privacy sensitive information. All of the data used for the simulation study will is maid available via Zenodo (see STAR methods).
- All original code has been deposited at GitHub and Zenodo and is publicly available (see STAR methods).
- Any additional information required to reanalyze the data reported in this paper is available from the lead contact upon request.

## Method Details

### List of DataSHIELD client-side functions

We list all functions that are used within the DataSHIELD framework for the client-side [Huth, 2023a] and explain their functionalities. The main function is `ds.did` which computes treatment effects and standard errors and calls the other client-side functions while being executed.

### Federated sample analogues for point estimates

In more detail, the computations are

1. 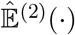: The client computes the sample mean of the variable *G*_*g*_ indicating if an individual has been treated in period *g* using a federated mean (`ds.mean` from dsBaseClient). The aggregate mean is then sent back to the servers (`ds.SendToServer`) for further analysis and processing.
2. 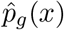: The client computes the parameter estimates (`ds.glm` from dsBaseClient), denoted by 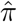, of a logistic regression model using a federated generalized linear model with the individuals that have been treated in period *g* or are never treated

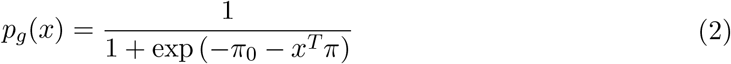 The estimated parameters, 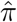, are then sent to the servers where they are immediately processed to compute the estimated probability of treatment, 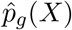, using (2) (`ds.genProp`). If no covariates are used, the probability of treatment can be directly computed using a federated mean of the variable indicating treatment *G*_*g*_ (`ds.mean`).
3. 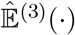: With the knowledge of the estimated probability of treatment, 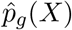, from step 2 and the binary variable indicating if the individual is part of the never-treated group, *C*, the mean of the expected value, 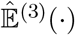, can be computed using a federated mean (`ds.mean`).
4. 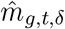: The client computes the parameter estimates 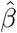 (`ds.glm`) of a linear regression model using the individuals that have never been treated and the ones that are treated in *g*. The linear regression model is represented by

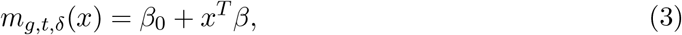

which relates the expected difference in the outcome of the control group and the covariates *X*. Subsequently, the client sends the estimated parameters 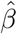 to the servers, where they are immediately processed to compute 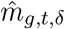 by (3) (`ds.multiplyMatrixMatrix`). If no covariates are used, the client can directly compute 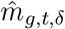 by taking the federated mean of *Y*_*t*_ − *Y*_*g*−*δ*−1_ (`ds.mean`).
5. 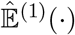: Finally, the client computes the expectation 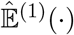 by using a federated mean (`ds.mean`) of all the known quantities and parameters. This expectation represents the point estimate of the average treatment effect of the treated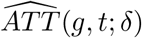.

### Federated computations of influence functions

The influence function, denoted as *ψ*_*g*,*t*,*δ*_(*W*_*i*_, *β*_*g*,*t*,*δ*_), measures the impact that each individual has on the estimated treatment effect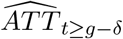. The collection of these influence functions 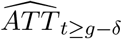 is then used to compute (approximate) the variance of 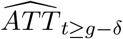 through the multiplier bootstrap method or asymptotic standard errors. In this section, we demonstrate the federation of the influence function for the DR estimate using the never-treated individuals as the control group. To maintain consistency with the notation used in [Callaway and Sant’Anna, 2021], we adhere closely to the notation used in that paper throughout this section. It is also worth noting that, following [Callaway and Sant’Anna, 2021], the parameters *β, π* used to estimate *m*_*g*,*t*,*δ*_(*X*), *p*_*g*_(*X*) are included in this section.

The theoretical influence function *ψ*_*g*,*t*,*δ*_(*W*_*i*_, *β*_*g*,*t*,*δ*_), as defined in [Callaway and Sant’Anna, 2021],

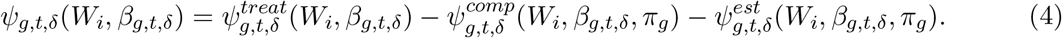

quantifies the effect of each individual on the treatment estimate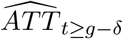. We will show in the following how this influence function can be estimated on the server sides in a federated manner using sample analogues, denoted by hat variables.

The components of the estimated influence function are given by [Callaway and Sant’Anna, 2021]

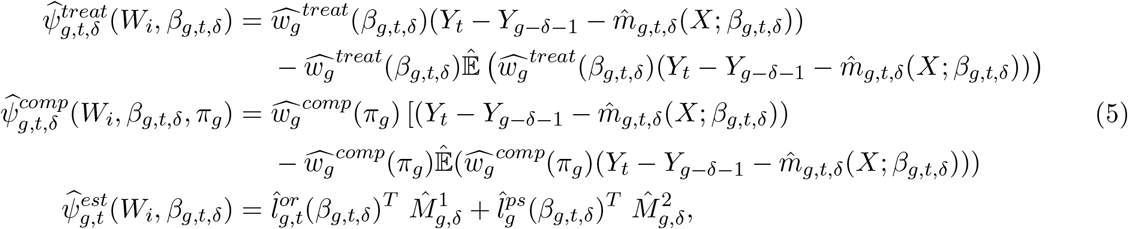

where *W*_*i*_ is the collection of all individual information about *Y, X*, and *C*. The weights are defined as 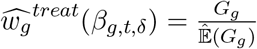 and 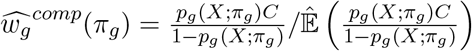 and the asymptotic linear representations of the regression estimator for the outcome evaluation of the comparison group and the generalised propensity score are

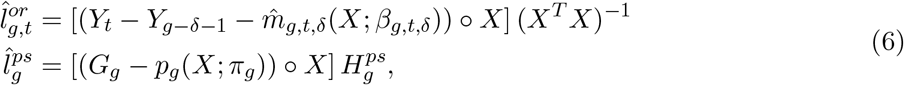

where *a* ° *B* defines a columnwise Hadarmad product between the vector *a* ∈ ℝ^*n*^ and the matrix *B* ∈ ℝ^*n×k*^. 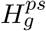 is the hessian obtained from the regression on *p*_*g*_ (*X*; *π*_*g*_).

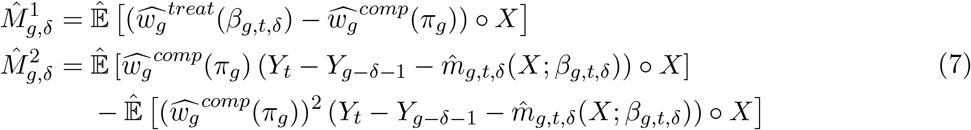

where *π*_*g*_ is the analogue of *π* in equation (2) and *β*_*g*,*t*,*δ*_ is the analogue of *β* in equation (3).

To compute the influence function in a federated manner, the sample analogues of the theoretical influence function must be computed on the client side. Specifically, the client must compute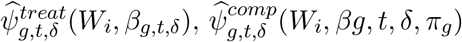, and 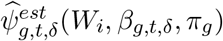 using the individual data available on the servers. These sample analogues will be used to estimate the noise-corrupted influence function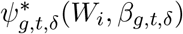.

1. 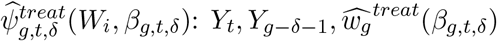, and 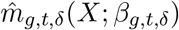 are known from the federated point estimate computations on the server side. 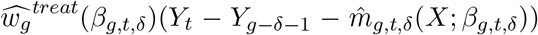 can therefore be computed on the client side (`ds.make` from dsBaseClient). Subsequently, the federated expectation can be returned to the client side (`ds.mean`) and be send back to the server side (`ds.SendToServer`). Finally, 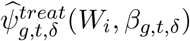 can be computed on the server side (`ds.make`).
2. 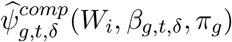: Analogously to 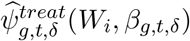 using 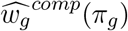 instead of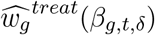.
3. 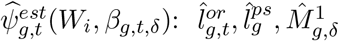, and 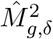 are unknown to the server side after the federated point estimate computations.
  a. 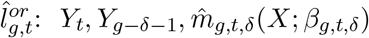, and *X* are known from the federated point estimate computations on the server side. (*X*^*T*^ *X*) can be computed via a federated aggregation (`ds.computeMatrixCrossproduct`) and be send to the client side, where its inverse (*X*^*T*^ *X*)^−1^ is computed and send to the servers. Subsequently, 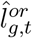 can be computed on the server side (`ds.multiplyMatrixMatrix`).
  b. 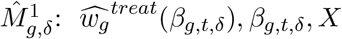 and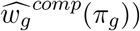, are known to the server side from the federated point estimate computations. Such that 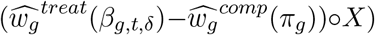 can be computed on the server side (`ds.make`) and the exp ectation can be estimated on the client side using a federated mean (`ds.mean`). The federated mean is subsequently send back to the servers (`ds.sendToServer`).
  c. 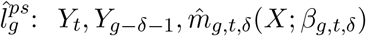, and *X* are known from the federated point estimate computations on the server side such that *G*_*g*_ − (*p*_*g*_(*X*; *π*_*g*_)) ° *X* can be computed on the server side (`ds.make`). 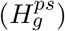 can be send to the server side where it is immediately multiplied to compute 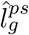 on the server side (`ds.multiplyMatrixMatrix`).
  d. 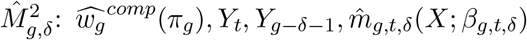 and *X*, are known to the server side from the federated point estimate computations. 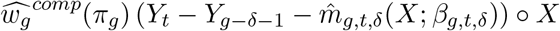 and 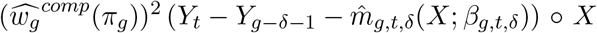 can therefore be computed on the server side (`ds.make`). The expectations can be returned to the client side using a federated mean (`ds.mean`). The means are subsequently send back to the servers (`ds.sendToServer`).

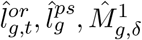 and 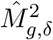 are now known to the server side. 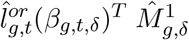 and 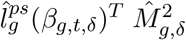 can therefore be computed on the servers (`ds.multiplyMatrixMatrix`). Subsequently, 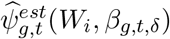 can be computed on the server sides (`ds.make`).

Knowing 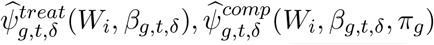 and 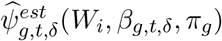 on the server sides, 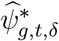 can be computed on the server sides (`ds.make`).

### Federated asymptotic variance-covariance matrix

In this subsection, we will demonstrate the estimation of the asymptotic variance-covariance matrix of the average treatment effect’s estimates in a federated setting.

The asymptotic distribution of the ATT is defined in [Callaway and Sant’Anna, 2021] as

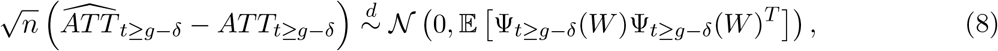

where *ATT*_*t*≥*g*−*δ*_ is the collection of *ATT* (*g, t*; *δ*) with *t* ≥ *g* − *δ* and Ψ_*t*≥*g*−*δ*_(*W*) is the collection of relevant influence functions *ψ*_*g*,*t*,*δ*_.

In order to estimate 𝔼 [ Ψ_*t*≥*g*−*δ*_(*W*)Ψ_*t*≥*g*−*δ*_(*W*)^*T*^ ], we use the sample analogue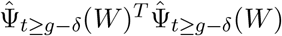, where the number of rows in the matrix 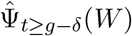 corresponds to the total number of individuals and each column represents a combination of treatment periods *g* and treatment evaluation periods *t*. The federated matrix product is computed using (`ds.computeMatrixCrossproduct`), which we explain more detail within the section Federated matrix product with its transpose.

### Federated multiplier bootstrap

The multiplier bootstrap allows computations of (clustered) standard errors. Callaway and Sant’Anna [2021] define one draw of the multiplier bootstrap as

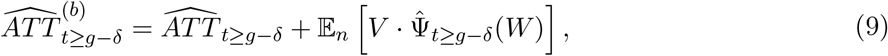

where 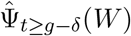 is the sample analogue of Ψ_*t*≥*g*−*δ*_(*W*) and 𝔼_*n*_(*a*) is the expectation over the components of a vector *a. V* is a random variable with 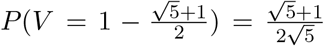 and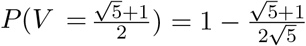. We subsequently show that the multiplier bootstrap can be federated using an example with *s* servers. By running the computations on each server (`ds.multiplierBootstrap`) and aggregating the results via a federated mean (`ds.mean`), the client can, due to the randomness of *V*, not exactly reproduce the results but obtain the same results qualitatively. Let 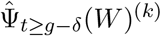 be the estimated influence matrix and *v*^(*k*)^ be the vector or observations drawn from *V* on server *k*, such that

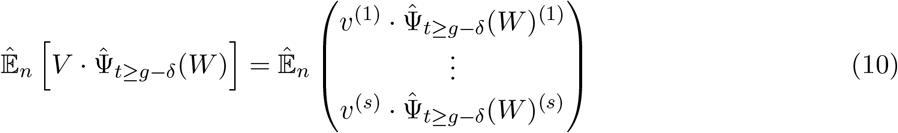

The multiplier bootstrap can be run on each server (`ds.multiplierBootstrap`) and multiplied by the server side sample size in order to compute each row of the matrix on the right hand-side of (10) on the respective servers. The client subsequently returns an estimate of the expectation using (`ds.mean`). The variance-covariance matrix of the average treatment effect’s estimate can finally be estimated with *B* bootstrap draws 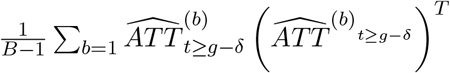.

### Federated matrix product with its transpose

To compute asymptotic standard errors, the matrix product of the transposed data and itself must be returned to the client side. This allows for the calculation of the asymptotic linear representation, denoted 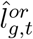, of the outcome regression in (7). In our package, this is done via the function `ds.computeMatrixCrossproduct`. Let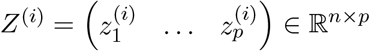, for *i* = 1, 2, …, *s*, be data matrices of *s* different servers such that the full data *Z* is given, without loss of generality, by

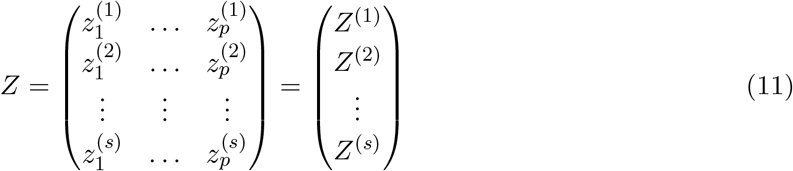

The matrix product of it’s transpose is therefore given by

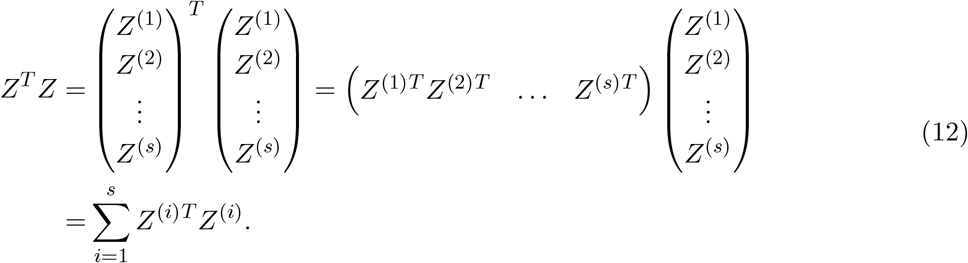

The multiplication of the server-side data with its transposed *Z*^(*i*)*T*^ *Z*^(*i*)^ can be computed on each server and returned to the client side if it contains at least as enough different entries as allowed by the data providers via DataSHIELDS disclosure settings (default is 5). Knowing all *Z*^(*i*)*T*^ *Z*^(*i*)^ on the client side, the sum can subsequently be computed on the client side to obtain *Z*^*T*^ *Z*.

### Federated column appending of influence functions

While iterating over the treatment periods *g* and the treatment impact periods *t*, the data frames are subsetted with respect to the individuals who are either treated in *g* or in the control group. On the client side, the influence function is computed for these subsetted individuals. To apply the clustered multiplier bootstrap, however, the influence function must be sent back to the server side after each iteration of *g* and *t*. Since the individuals change every iteration, simply appending the influences column-wise is not possible. Therefore, we create a data frame filled with zeros and one column indicating the ID of the individuals, which is known to the server. However, appending a full vector, even though the vector is stored on ther server side, poses a data security problem as it allows for the Covariance-Based Attack Algorithm described in [Huth et al., 2023]. To address this, the rows of the data frame are randomly shuffled after each iteration.

To enhance understanding, we present a simplified example utilizing 4 individuals, two treatment periods *g* = 1, 2, and two treatment impact periods *t* = 3, 4. We assume that individual id_1_ is treated in *g* = 1, individual id_3_ is treated in *g* = 2, and individuals id_2_ and id_4_ are never treated. The initialized data frame ℱ_0_ would be given by

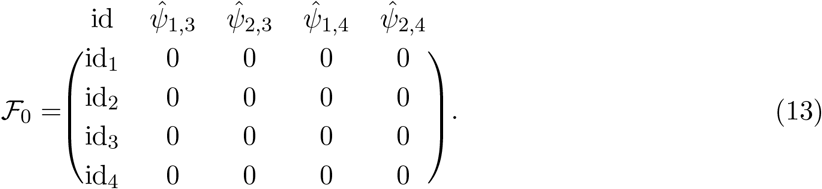

At the first iteration, the server stores the ID vector *v*_1,3_ = (id1 id2 id4)^*T*^ of the individuals used in this run. The server computes the influence vector *w* _1,3_ = (3 −1 2)^*T*^. On the server, the ID vector and the influence vector are matched component-wise to create the matrix (*v*_1,3_ *w*_1,3_). In the next step, the relevant IDs in the first column of ℱ_0_ are replaced with the corresponding values from *w*_1,3_, and the matrix is subsequently shuffled row-wise

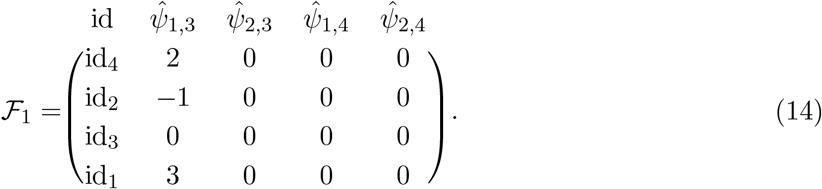

This procedure is repeated for all combinations of *t* and *g* such that the final data frame is, depending on the random shuffling, given by

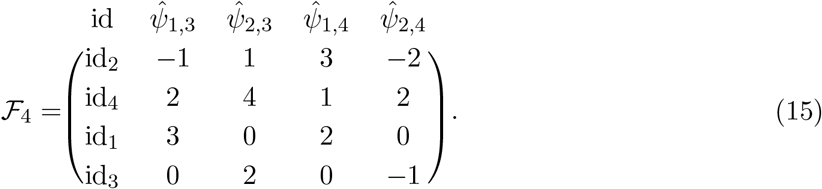

By implementing this method, it is possible to append the influence functions in a privacy-preserving manner.

### Summary statistics - Malaria dataset

## Supplementary Material

